# Monoclonal antibody-based serotyping of Listeria monocytogenes provides new insights in epidemiology and virulence

**DOI:** 10.64898/2026.05.20.726485

**Authors:** Jasper Mol, Kim Duindam, A. Robin Temming, Rob van Dalen, Yvonne Pannekoek, Nina M. van Sorge

## Abstract

**Objectives:** *Listeria monocytogenes* is an opportunistic pathogen, associated with foodborne infections that disproportionately affect newborns, elderly and immunocompromised patients. *L. monocytogenes* can be classified on the antigenic and related structural variation of cell-associated wall teichoic acid (WTA) molecules through conventional serotyping techniques. The WTA structure of serovars (SV) 1/2, 1/2*, 3 and 7 consists of a linear poly-ribitolphosphate (RboP) polymer either with or without decoration with rhamnose (Rha) and/or N-acetylglucosamine (GlcNAc). Of these four SVs, SV1/2 (WTA with GlcNAc and Rha) causes ∼ 99% of all listeriosis cases. However, conventional serotyping cannot accurately discriminate between these four SVs, particularly SVs1/2* (WTA with Rha).

**Methods:** Here we applied two identified monoclonal antibodies (mAb), with specificity for the RboP backbone or GlcNAc modification to develop a discriminatory serotyping scheme for SV1/2, 1/2*, 3 and 7. Isogenic mutants for the different SVs were created in *L. monocytogenes* SV1/2 strain EGD-e. The typing scheme was then adapted to an immnoblot assay and applied to a collection of 317 previously classified listeriosis isolates from the Netherlands Reference Laboratory for Bacterial Meningitis.

**Results:** Binding of the RboP-specific mAb was limited to EGD-e wild type (SV1/2), but increased significantly for isogenic EGD-e mutants representing SV1/2*, 3 and 7. In contrast, the GlcNAc-specific mAb only recognized EGD-e mutants representing SVs 1/2 and 3. The combined staining profiles of the two mAbs allowed accurate discrimination of the four SVs as verified on clinical isolates. Applying this typing scheme to 317 listeriosis isolates previously typed as SV1/2, we confirmed SV designation in >90% of isolates, but also identified SV1/2* (5.4%), SV3 (0.6%) and SV7 (0.3%) isolates. SV1/2* isolates were also identified among meningitis patients.

**Conclusion:** The increased discriminatory capacity of *L. monocytogenes* serotyping provides a more detailed insight of the epidemiological landscape and the critical factors for *L. monocytogenes* infections.

## Introduction

*Listeria monocytogenes* is a Gram-positive, intracellular, opportunistic pathogen. The route of infection for *L. monocytogenes* is often through contaminated foods, such as soft cheeses smoked seafood or processed deli meats. Newborns, pregnant women, elderly people and immunocompromised patients are most at risk for listeriosis and results in a near-100% hospitalization rate with a mortality of 20-25% (1). Furthermore, listeriosis is on the rise, with a peak incidence in Europe in 2024, highlighting the necessity for adequate surveillance and food safety regulations (2).

*L. monocytogenes* can be classified into 13 serovars (SVs), which can be serologically differentiated based on structural and compositional variation of wall teichoic acid (WTA), also referred to as the somatic (O) antigen, and the flagellar (H) antigen. *L. monocytogenes* WTA consists of ribitolphosphate (RboP) polymers that are covalently anchored to the peptidoglycan layer of the bacterial cell wall. *L. monocytogenes* WTA can be broadly divided into two structural variations. Type I WTA consists of repeating linear RboP subunits, whereas the repeating unit of type II WTA contains an N-acetyl-glucosamine (GlcNAc) residue on the C-2 or C-4 positions of RboP (3). Both WTA types can be further modified through glycosylation, giving rise to specific SVs. For type I WTA, four SVs are described: SV1/2, 1/2*, 3 and 7 (Figure 1A). SV1/2 is decorated with *N*-acetylglucosamine (GlcNAc) at the C-2 position and rhamnose at the C-4 position of the RboP monomers. SV1/2* and SV3 are variants of SV1/2, missing the GlcNAc and rhamnose moieties, respectively. Finally, SV7 expresses unglycosylated RboP WTA. From bacteriological surveillance data, which relies on classical serotyping methods, SV1/2 accounts for approximately one-third of all invasive infections in the Netherlands (4).

**Figure 1.**
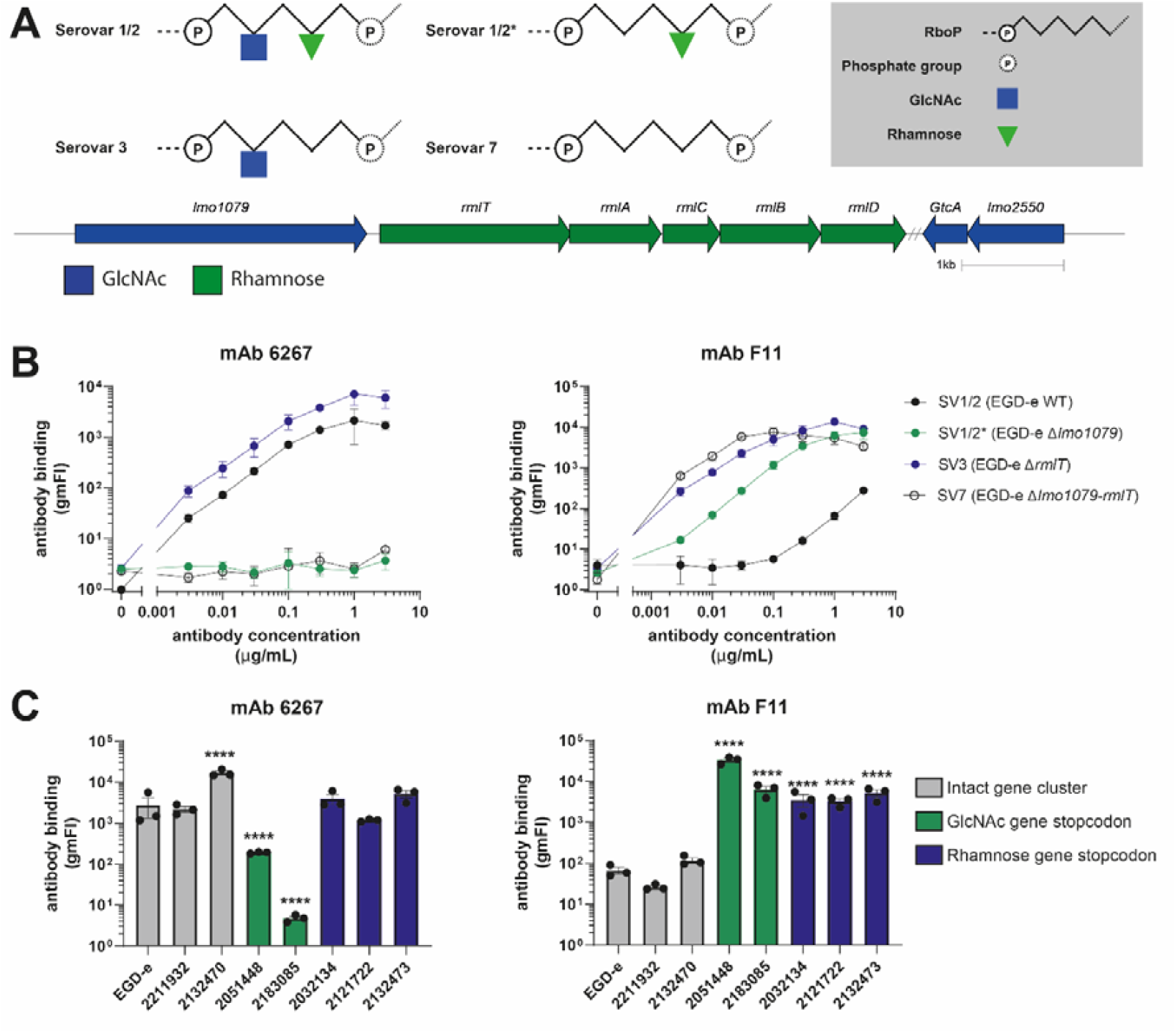
*L. monocytogenes* type I WTA serovars are discriminated with specific monoclonal antibodies. **A**. The schematic representation of type I WTA consisting of a linear RboP polymer backbone with further glycosylation by GlcNAc and rhamnose. The presence of these WTA glycosylations is conferred by specific genes, i.e. *lmo*1079 for GlcNAc and *rmlT* for rhamnose. **B**. SV-specific antibody binding profile of 6267 and F11 to isogenic type I WTA mutants in EGD-e background. Binding is shown as the geometric mean of fluorescence intensity of three biological replicates ± SEM. **C**. mAb 6267 and F11 (both at 1 µg/mL) binding to clinical *L. monocytogenes* isolates. Whole genome sequencing data was analyzed for the presence or absence of premature stopcodons in genes related to WTA GlcNAcylation or rhamnosylation. Antibody binding to these isolates was compared to EGD-e wild-type with a log-normal One-Way-ANOVA. ****: p<0.0001.

WTA structural variability results in different immunogenicity, allowing for the development of *L. monocytogenes* serotyping methods. Serotyping was originally developed as an agglutination technique using polyclonal antisera (5). However, a multicenter study concluded that the reproducibility of this agglutination assay is highly variable, highlighting the critical need for high quality serotyping materials (6). Furthermore, SV1/2* cannot be discriminated with these antisera.

PCR assays have been developed for SV determination, however these do not discriminate between the four Type I WTA SVs (7, 8). Among these, only SV1/2 is regarded as clinically relevant (9). However, this classification is based on the previously mentioned serotyping methods that are incapable of reliably distinguishing between type I WTA SVs.

These mentioned limitations in *L. monocytogenes* serotyping methodologies may have resulted in a skewed epidemiological view of listeriosis. As an alternative to the serotyping assays, Sumrall and colleagues developed a method based on the binding of bacteriophage proteins to specific WTA glycan motifs (10). This improved *L. monocytogenes* SV discrimination compared to conventional techniques. However, this technique relies on access to specific laboratory equipment to analyze and quantify the read out, i.e. fluorescence. As the global burden of listeriosis disproportionately affects developing regions, this equipment may not be readily available (11). To this end, we have developed a high throughput and accessible assay that can accurately discriminate the four type I WTA SVs, i.e. SV1/2, SV1/2*, SV3 and SV7. We used monoclonal antibodies (mAbs), in contrast to the conventional use of polyclonal antisera, and applied these to a colony immunoblot protocol that we previously used to assess WTA glycosylation in *Staphylococcus aureus* (12). We created an isogenic strain set representing the four type I WTA SVs to demonstrate SV-specific binding profiles of the mAbs, first with flow cytometry and subsequently with an immunoblot protocol. Finally, we applied this approach to a collection of clinical *L. monocytogenes* isolates from the Netherlands Reference Laboratory for Bacterial Meningitis (NRLBM) that were previously serotyped as SV1/2 with commercial antisera to determine the presence of non-SV1/2 strains.

## Materials and Methods

### Bacterial strains and culturing conditions

All bacterial strains used in this study are listed in supplementary table S1. *L. monocytogenes* strains were routinely grown overnight at 37°C on Brain Heart Infusion (BHI; Oxoid) or Columbia blood agar plates (Oxoid), or in BHI broth with agitation, supplemented with antibiotics when required. *Escherichia coli* strains were cultured at 37°C on Luria-Bertani (LB; Oxoid) plates or in LB broth with agitation.

### Monoclonal antibody production

Monoclonal antibodies used in this study were produced as described previously (13). In short, mammalian expression plasmids (pcDNA3.4) encoding the human IgG1 heavy chain and human kappa light chain sequences were transfected 1:1 into human embryonic kidney (HEK) 293 Freestyle cells using Polyethylenimine MAX® (PEI MAX) as transfection reagent. After 5 days of shaking at 37°C and 8% CO_2_, supernatants were harvested by centrifugation and filtered through a 0.2 um filter. Subsequently, protein G agarose (Pierce) was added to the supernatants and incubated overnight at 4°C on a tube roller. Protein G-bound human IgG1 mAbs were consecutively washed with phosphate-buffered saline (PBS), and eluted with 0.1 M Glycine pH 2.5. The eluted fraction was directly neutralized using 1 M Tris pH 8.7 and buffer-exchanged to PBS through centrifugation in a Vivaspin concentration tube (Sartorius, 100 kDa MWCO).

### *L. monocytogenes* mutagenesis

All primers and plasmids used in this study are listed in supplementary table S2. Deletion of the genes encoding the glycosyltransferases essential for GlcNAc- (*lmo*1079) or rhamnose decoration (*rmlT*) was performed using the pMAD plasmid, as described previously (14). A detailed description is found in the supplementary materials.

In short, the up- and downstream regions of each gene were amplified and cloned into the pMAD vector using restriction enzymes. These plasmids were transformed into competent EGD-e WT cells and transformants were selected at 30ºC using erythromycin resistance and a colour change in colonies from white to blue. Successful transformants were incubated under erythromycin pressure at 43ºC for plasmid integration. Resistant colonies were grown overnight at 30ºC without antibiotic pressure and colonies that had lost erythromycin resistance and the blue colour were checked for gene loss.

### Monoclonal antibody binding assay to *L. monocytogenes*

L. *monocytogenes* overnight cultures were diluted 1:20 in fresh BHI and grown to mid-exponential phase (∼OD_600_ = 0.5). Cells were harvested by centrifugation (3,000 x g, 4ºC, 10 minutes) and resuspended in PBS supplemented with 0.1% Bovine Serum Albumin (Sigma Aldrich) (PBS + 0.1% BSA). The bacterial suspension was diluted in fresh PBS + 0.1% BSA to a concentration of approximately 10^8^ colony forming units (CFU)/mL. Antibodies were added to 10^6^ CFU bacteria across a concentration range (0-3 µg/mL) or at a specific concentration in a final volume of 25 µL in a 96-wells plate and incubated statically at 4ºC for 20 minutes. Bacteria were washed with PBS + 0.1% BSA and subsequently incubated with 25 µL Human α-kappa AF647 (Southern Biotech; 1:200 in PBS +0.1 % BSA) statically at 4ºC for 20 minutes. Finally, the bacteria were washed once more with PBS 0.1% BSA and fixed with PBS + 1% formaldehyde (Sigma Aldrich) and fluorescence intensity was measured on FACSCanto or FACSymphony (BD Biosciences). Flow cytometry data was subsequently analyzed using FlowJo (BD Biosystems).

### Colony immunoblot

For each *L. monocytogenes* strain, a single overnight colony was resuspended in 50 µL PBS and 3 µL droplets were spotted and grown overnight on Colombia blood agar plates. The next day, colonies were transferred to a nitrocellulose membrane and air-dried for 20 minutes. All washing and staining steps were performed in 50 mL tubes (Falcon) on a tuberoller (35 rpm). The membrane was first washed three times with demineralized water for five minutes and twice with Tris Buffered Saline (TBS; Roche). Next, the membrane was blocked with TBS + 0.1% Tween-20 (TBST; Thermo) + 5% BSA for one hour, after which the membrane was incubated with either monoclonal antibody 6267 or F11 (final concentration 0.5 µg/mL in TBST + 2.5% BSA) for one hour. After washing three times with TBST (5 minutes each), the membrane was incubated for one hour with Goat F(ab’)2 Anti-Human Kappa-AP (Southern Biotech; 1:5,000 dilution in TBST + 2.5% BSA). After washing three times with TBST, antibody binding was visualized using the Vector blue AP Substrate kit according to the manufacturer’s instructions (VectorLabs).

### Statistical analysis

Data are presented as the geometric mean of fluorescence intensity ± standard error of mean (SEM). Statistical analyses were performed using Graphpad Prism 10.6 (GraphPad Software). Antibody binding data obtained at specific concentrations were analyzed by one-way analysis of variance (ANOVA) followed by Bonferoni’s multiple comparison test. *P* < 0.05 was considered as significant difference in binding.

## Results

### WTA-specific mAbs bind to distinct glycan profiles present on *L. monocytogenes*

Conventional serotyping techniques lack complete and reliable discrimination between the four type I WTA-producing *L. monocytogenes* isolates. Therefore, we aimed to develop an assay to address this limitation. As the WTA glycosylation motif determines whether the strain is typed as SV1/2, SV1/2*, SV3 or SV7 (Figure 1A), we aimed to use monoclonal antibodies that could bind to specific WTA glycosylation profiles. To develop this approach, we first created a set of isogenic deletion mutants for the various type I WTAs, using the wildtype SV1/2 strain EGD-e. By deleting the glycosyltransferase essential for GlcNAc decoration (*lmo*1079, Figure 1A) and for rhamnose (*rmlT*, Figure 1A), we created EGD-e-Δ*lmo*1079, and EGD-e Δ*rmlT*, which lacks GlcNAc (SV1/2*) and rhamnose (SV3), respectively. Finally, we deleted both genes in the same strain (EGD-e Δ*lmo*1079*-rmlT*), resulting in an SV7 phenotype.

Next, we assessed binding of two mAbs to this isogenic set of EGD-e strains. mAb 6267 was originally isolated and characterized based on recognition of *S. aureus* WTA decorated with α-GlcNAc (15) and showed binding to both wildtype *L. monocytogenes* EGD-e bacteria (SV1/2) and EGD-e Δ*rmlT* (SV3) in a concentration and saturable manner (Figure 1B). In contrast, deletion of lmo1079, resulting in loss of GlcNAc on WTA (SV1/2*), and deletion of both Δ*lmo*1079-*rmlT* (SV7), leaving the unglycosylated RboP backbone, resulted in complete loss of 6267 binding. Thus, 6267 specifically recognizes α-GlcNAc on *L. monocytogenes* WTA.

Monoclonal antibody F11 was identified and characterized in-house as recognizing the WTA RboP backbone (13). We assessed binding of F11 to the same panel of isogenic *L. monocytogenes* strains. Similar to 6267, F11 bound to the Δ *rmlT* mutant (SV3) (Figure 1B). However, in contrast to 6267, F11 also bound to the Δ*lmo*1079 (SV1/2*) and Δ*lmo*1079-*rmlT* (SV7), but not to the wildtype bacteria (SV1/2). These results indicate that binding of F11 is blocked by the presence of the combination of GlcNAc and rhamnose residues on SV1/2 WTA, but binding occurs upon the loss of either modification.

The binding profiles of 6267 and F11 to the EGD-e wild-type and isogenic deletion mutants allowed for discrimination between *L. monocytogenes* strains belonging to SVs 1/2, 1/2*, 3 and 7. To this end, we examined binding of these antibodies to clinical listeriosis isolates, that were previously collected and typed by the Netherlands Reference Laboratory for Bacterial Meningitis (NRLBM) as SV1/2 or SV3, and for which genome sequences were available. Two isolates, 2211932 and 2132470, showed binding to 6267, but not to F11, similar to the EGD-e WT strain. Correspondingly, the genes encoding the GlcNAc and rhamnose biosynthesis pathways were intact in these strains. In contrast, the clinical isolates 2051448 and 2183085 showed reduced binding to 6267 but increased binding to F11, when compared to EGD-e. Correspondingly, both strains had a premature stopcodon in one of the WTA-GlcNAc biosynthesis genes, indicating that these isolates have lost WTA-GlcNAc decoration. Lastly, isolates 2032134, 2121722 and 2132473, showed the same binding to 6267 but increased binding to F11 compared to EGD-e, and genes associated with WTA rhamnosylation contained premature stopcodons, suggesting that the WTA-Rha was lacking.

Overall, the specific binding patterns of mAbs 6267 and F11 allowed for accurate discrimination of the four different type I WTA SVs. In case no binding of mAb 6267 or F11 is observed, this indicates that the *L. monocytogenes* expresses type II WTA and most likely belongs to SV4b (Supplemental figure 1).

### Colony blot as an accessible, high-throughput platform to discriminate type I WTA-producing *L. monocytogenes*

We next aimed to translate the discriminatory characteristics of our mAbs into an accessible and high-throughput assay that can be applied in any laboratory setting. To this end, we transformed our flow cytometry-based assay into an immunoblot protocol previously used to detect specific WTA glycopatterns in *S. aureus* (Figure 2A) (12). In this assay, *L. monocytogenes* strains grown on agar plates are transferred to duplicate nitrocellulose membranes and stained with the respective mAbs. After development of the blot, antibody binding results in a blue colored colony. Since 48 colonies can be spotted onto a single agar plate, many isolates can be serotyped in a single experiment.

**Figure 2.**
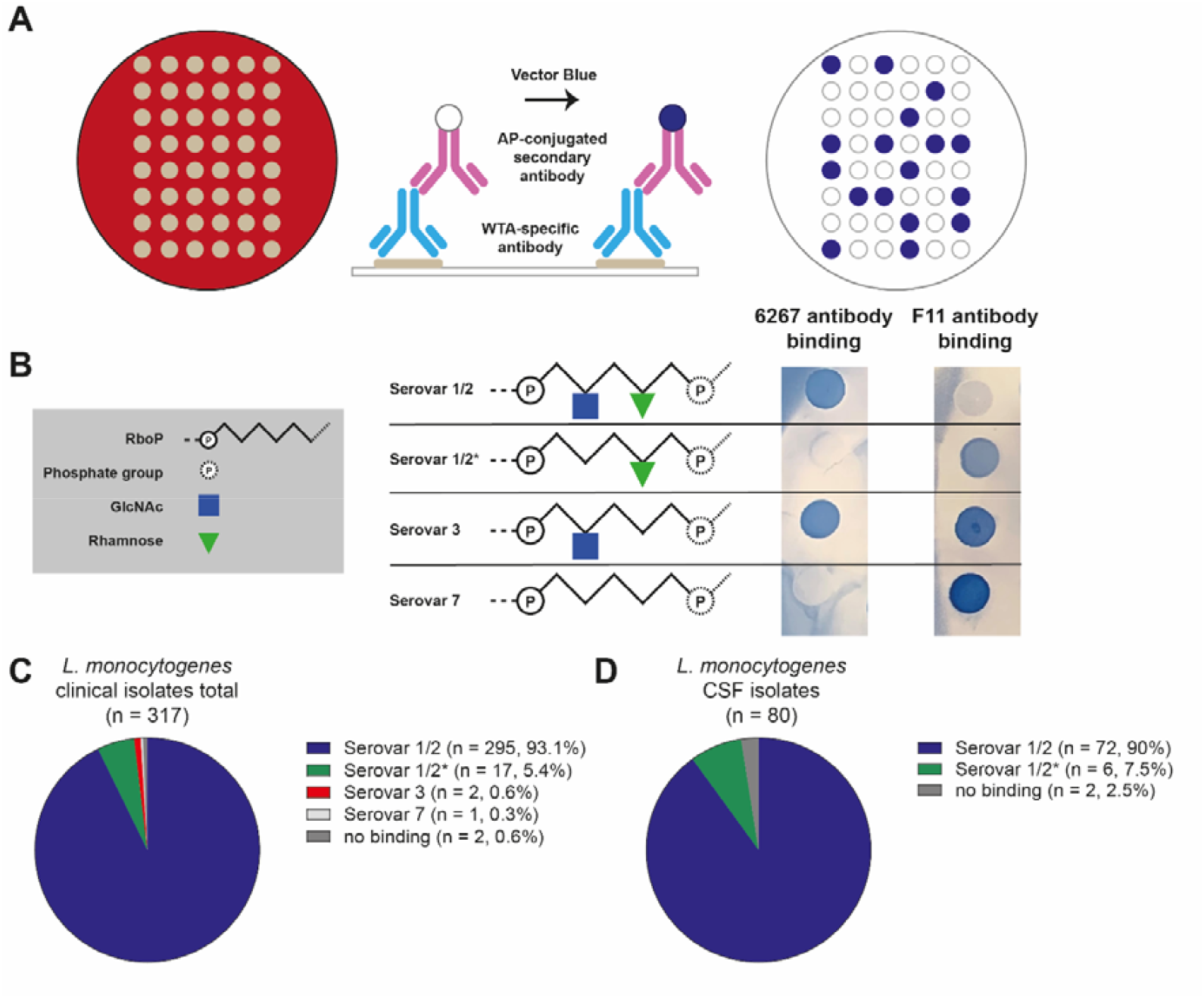
Monoclonal antibody-based immunoblot serotyping of clinical *L. monocytogenes* isolates. **A**. *L. monocytogenes* isolates are spotted on an agar plate and grown overnight. The colonies are transferred to duplicate nitrocellulose membranes and stained with WTA-specific monoclonal antibodies 6267 and F11. **B**. Immunoblot readout is based on the binding of antibodies 6267 and F11. Shown are the binding of these antibodies to EGD-e WT (serovar 1/2) and isogenic glycosyltransferase mutants (corresponding to SV1/2*, 3 and 7). **C**. SV distribution of 317 clinical isolates based on the immunoblot assay. **D**. SV distribution of 80 *L. monocytogenes* strains isolated from the CSF of patients with bacterial meningitis.

We first validated this method on our set of EGD-e isogenic SV variants (Figure 2B). The binding pattern on the immunoblots overlapped with flow cytometry results, with antibody 6267 binding to both the SV1/2 and the SV3 strains, and the F11 antibody binding to SV1/2*, SV3 and SV7 strains. Interestingly, for the F11 antibody, we consistently observed a more pronounced color development for SV7 compared to SV1/2*, suggestive of a higher deposition of F11 on SV7 WTA. This indicates that color intensity may discriminate the two SVs.

### Immunoblot serotyping of clinical *L. monocytogenes* isolates from the Netherlands

Next, we applied our immunoblot approach to *L. monocytogenes* isolates that had been collected and typed by the NRLBM as SV1/2 using commercially available typing sera. Out of the 317 isolates, 295 (93.1%) were confirmed as SV1/2 based on the immunoblot method (Figure 2C). However, we also identified 17 (5.4%) SV1/2* isolates, two (0.6%) SV3 isolates, and one (0.3%) SV7 isolate. Finally, we identified two *L. monocytogenes* strains that did not show binding to either antibody, suggesting that these isolates do not express type I WTA.

In some patients, *L. monocytogenes* can cross the blood-brain barrier to cause meningitis. To assess whether there is a different SV distribution in patients with or without meningitis, we plotted serotyping results for 80 isolates cultured from cerebrospinal fluid (CSF), the hallmark of meningitis. Of these 80 isolates, 72 (90%) isolates were SV1/2 for, and six (7.5%) isolates were SV1/2* (Figure 2C). Two isolates were negative for both antibodies in the blot, suggesting a type II WTA SV (most likely SV4b). Overall, the SV distribution was similar for *L. monocytogenes* strains from meningitis patients compared to all listeriosis patients.

## Discussion

L. *monocytogenes* strains can be divided into 13 SVs, based on the chemical composition of their WTA. For complete epidemiological analysis, an accurate serotyping approach is crucial. Conventional serotyping techniques have either been inconsistent or incomplete in their ability to accurately discriminate between different *L. monocytogenes* SVs. Previously, bacteriophage proteins that recognized SV-specific WTA structures were used for accurately discrimination, showing the potential of a glycan-based serotyping approach (10). Here we present an immunoblot serotyping assay for type I WTA *L. monocytogenes* bacteria based on WTA-specific monoclonal antibodies. This approach is high-throughput and accessible, as it does not require advanced equipment or computational analysis. We validated our assay using isogenic SV knockout mutants and showed its reproducibility on clinical *L. monocytogenes* isolates that produce WTA corresponding to SV1/2, SV1/2*, SV3 and SV7.

We used our monoclonal antibody-based immunoblot assay to serotype 317 clinical *L. monocytogenes* isolates collected by the NRLBM that were previously serotyped as SV1/2 using the commercial antisera agglutination kit (Denka Seiken, Tokyo, Japan). Although the majority of isolates was serotyped as SV1/2 by the blot as well, a subset (5.4%) was serotyped as SV1/2*. Additionally, we identified two SV3 isolates and one SV7 isolate. Furthermore, among a subset of isolates obtained from the CSF of patients with bacterial meningitis previously typed as SV1/2, we found a comparable proportion of SV1/2* strains (6.9%). These results suggest that SV1/2* *L. monocytogenes* strains are capable of causing severe disease, including meningitis, which had not been recognized thus far. It would be interesting to extend this immunoblot assay to non-clinical isolates, obtained from food and environmental sources, since isolates from these sources often rely on PCR- or WGS serotyping, and SV1/2* would most likely not be identified consistently. Re-assessing the SV distribution in isolates from these sources would reveal whether the proportion of SV1/2* clinical isolates are over- or underrepresented in invasive human infection isolates (16).

The similar proportion of SV1/2* isolates in the total number of clinical *L. monocytogenes* isolates screened and the isolates from patient CSF suggest that the loss of GlcNAc on the WTA might not be reducing the ability of SV1/2* isolates to cross the blood-brain barrier. This SV maintains the presence of rhamnose on its WTA, which was shown to protect against killing by antimicrobial peptides and promote virulence in a mouse infection model (17). As *L. monocytogenes* infections have a high mortality rate, clinical patient cohort analyses of *L. monocytogenes* meningitis patients are routinely conducted (18-20). Unfortunately, we do not have sufficient patient information related to all our isolates to ascertain whether patients infected with SV1/2* have a different clinical profile compared to patients infected with SV1/2 strains. Extending our antibody-based immunoblot assay to other sets of clinical *L. monocytogenes* isolates would provide further insight into the SV distribution of *L. monocytogenes* and the virulence associated with specific SVs.

Further development of this serotyping application for *L. monocytogenes* would include antibodies specific for the various type II WTA SVs. Similar to type I WTA, SV differentiation in type II WTA is based on the glycosylation of the WTA backbone and changes are often the result of pointmutations in glycosyltransferases-encoding genes (21). Similarly to the F11 antibody used here, synthetic type II *L. monocytogenes* WTA fragments could be used to identify responsive B cell clones from human serum (13).

We developed a mAb-based serotyping assay that allows for increased discriminatory capacity of *L. monocytogenes* SVs and provides a more detailed insight of the epidemiological landscape. This assay can easily be expanded with additional antibodies, without the need for advanced equipment of computational analyses, making it an accessible approach for any reference or academic laboratory that regularly performs *L. monocytogenes* serotyping.

## Acknowledgements

The authors would like to thank Prof. Dr. Sven Halbedel for providing the *L. monocytogenes* EGD-e strain and the pMAD plasmid. Furthermore, they would like to thank the Netherlands Reference Laboratory for Bacterial Meningitis for their help with the selection of the relevant *L. monocytogenes* clinical isolates.

## Data availability statement

The sequences and materials for antibody F11 are currently not publicly available due to their inclusion in a related patent application (EP25187749.4, submitted July 2025; F11 is described as W2G2.1Uni). Nevertheless, the F11 antibody and its sequence will be made available for non-commercial research purposes upon request, subject to a Material Transfer Agreement (MTA) to protect intellectual property rights. Access to these materials can be requested by contacting Prof. dr. Nina M. van Sorge. These access restrictions are necessary due to ongoing intellectual property protections but do not preclude academic research use.

