## Supplemental information for "Monoclonal antibody-based serotyping of Listeria monocytogenes provides new insights in epidemiology and virulence"

Monoclonal antibody-based serotyping of Type I WTA expressing *Listeria monocytogenes*

Supplemental materials

*Corresponding author contact:

Professor Dr. Nina M. van Sorge

Amsterdam UMC, location AMC

Meibergdreef 9

IWO building, room IA3-211

1105 AZ Amsterdam, the Netherlands

**Supplemental methods**

For deletion of *lmo*1079 the up- (784 bp) and downstream (762 bp) regions were amplified from *L. monocytogenes* SV1/2 strain EGD-e chromosomal DNA with primers JM01+JM02 and JM03+JM04 and digested with BamHI/SalI and SalI/NcoI respectively. The digested products were cloned into BamHI/NcoI digested pMAD, creating pMAD-Δ*lmo*1079*.*

For deletion of *rmlT* the up- (940 bp) and downstream (831 bp) regions were amplified from EGD-e chromosomal DNA with primers JM07+JM08 and JM09+JM10. These products were merged using overlap-extension PCR and digested with MluI/NcoI and cloned into BamHI/NcoI digested pMAD, creating pMAD-Δ*rmlT*.

For deletion both *lmo*1079 and *rmlT,* the up- (784 bp) and downstream (1009 bp) regions were amplified from EGD-e chromosomal DNA with primers JM01+JM02 and JM13+JM14 and digested with BamHI/SalI and SalI/NcoI, respectively. The digested products were cloned into BamHI/NcoI digested pMAD, creating pMAD-Δ*lmo*1079*-rmlT*.

The resulting knockout plasmids (pMAD-Δ*lmo*1079, pMAD-Δ*rmlT* and pMAD-Δ*lmo*1079*-rmlT*) were transformed into *E. coli* DC10B and selected on ampicillin (100 µg/mL). Plasmid DNA was isolated using Miniprep (Genejet) to obtain adequate amounts of plasmid. Subsequently, these plasmids were electroporated into *L. monocytogenes* EGD-e, as described previously (1). *L. monocytogenes* transformants were selected on BHI supplemented with erythromycin (5 µg/mL) and 5-bromo-4-chloro-3-indolyl-β-D-galactopyranoside (X-gal) (50 µg/mL) (BHI + EX) at 30°C. Blue, erythromycin-resistant transformants were grown on BHI + EX at 42°C for 48 hours to obtain plasmid integrants. Integrants were grown overnight in BHI at 30°C, serially diluted and plated on BHI supplemented with 50 µg/mL X-gal (BHI + X). White colonies, indicating loss of plasmid, were plated out on both BHI + EX and BHI + X. White, erythromycin sensitive colonies were checked for the loss of gene with PCR and Sanger sequencing (primers JM05+JM06 for *lmo*1079 and JM11+JM12 for *rmlT*) and designated EGD-e Δ*lmo*1079*;* EGD-e Δ*rmlT* and EGD-E Δ*lmo*1079*-rmlT*.

**Supplemental table S1. Bacterial strains used in this study**

|  | **Source** |
| --- | --- |
| ***E. coli*** |  |
| DC10B | LMBP 9585 (2) |
| ***L. monocytogenes*** |  |
| EGD-e | (3) |
| EGD-e Δ*lmo1079* | This study |
| EGD-e Δ*rmlT* | This study |
| EGD-e Δ*lmo1079-rmlT* | This study |
| 2211932 | Netherlands Reference Laboratory for Bacterial Meningitis (NRLBM) |
| 2132470 | NRLBM |
| 2051448 | NRLBM |
| 2183085 | NRLBM |
| 2032134 | NRLBM |
| 2121722 | NRLBM |
| 2132473 | NRLBM |

**Supplemental table 2. Primers and plasmids used in this study.**

| **Primer** | **Sequence 5'-> 3'** |
| --- | --- |
| JM01 | AGTCGGATCCGGAGCATCTTCTACATTAGGC |
| JM02 | AGTCGTCGACCCATTAACTTTCTCCCTCC |
| JM03 | AGTCGTCGACTAAATGAGGGAAAACGTTAGG |
| JM04 | AGTCCCATGGCACCGTGAATGAACGCC |
| JM05 | GCAAATTGGAATGGGAGGCG |
| JM06 | GGATGCCTTGTTGCCGAAAC |
| JM07 | TTATCCATGGCCTAAAGTTAATGGCAAAGCTCCTGC |
| JM08 | CTTTTCTCTCCATTCTTAAAGCGCTCATTATATCCTCCTAAAATAG |
| JM09 | CTCATTATATCCTCCTAAAATAGCGCTTTAAGAATGGAGAGAAAAG |
| JM10 | TAATACGCGTCAACTGCAGCCTCATCAATAT |
| JM11 | TATTGCCACACGCTTTACCG |
| JM12 | CTTCCACGATTGAACGAACG |
| JM13 | CGGGTCGACTAAGAATGGAGAGAAAAGAATGAAAGG |
| JM14 | CGGCCATGGGGAATGCTTTTTCATTATAGC |
| **Plasmid** | **Source** |
| pMAD | (4) |
| pMAD-Δ*lmo1079* | This study |
| pMAD-Δ*lmo1080* | This study |
| pMAD-Δ*lmo1079-1080* | This study |

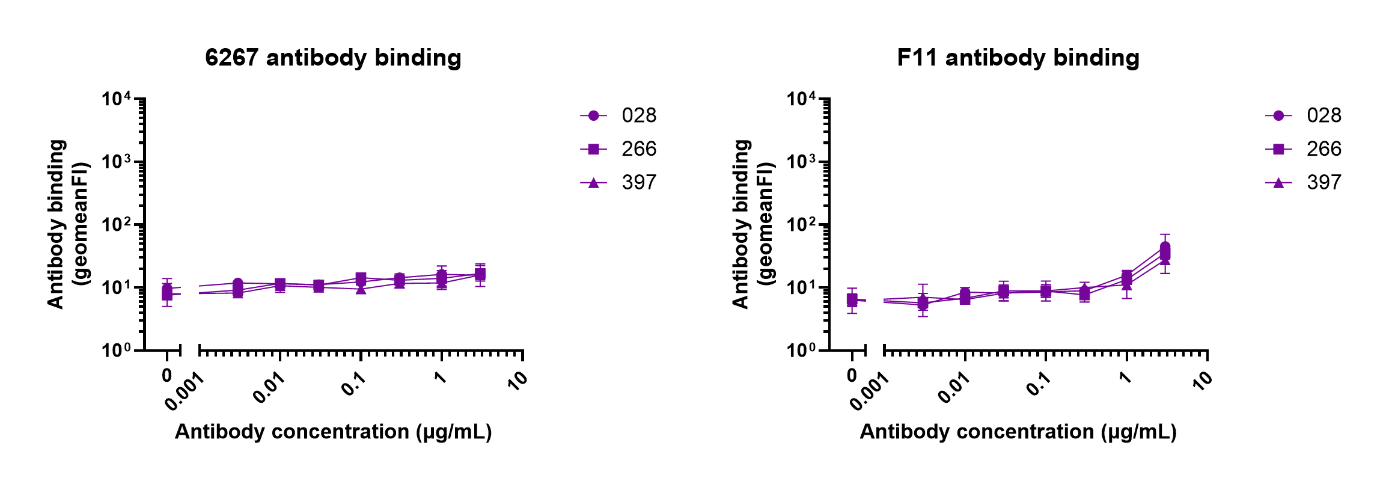

**Supplemental figure 1. *L. monocytogenes* serovar 4b isolates are not bound by antibody 6267 or F11.** Binding is shown as the geometric mean of fluorescence intensity of three biological replicates ± SEM.
